# DeepMotifSyn: a deep learning approach to synthesize heterodimeric DNA motifs

**DOI:** 10.1101/2021.02.22.432257

**Authors:** Jiecong Lin, Lei Huang, Xingjian Chen, Shixiong Zhang, Ka-Chun Wong

## Abstract

**Motivation:** The cooperativity of transcription factors (TFs) is a widespread phenomenon in the gene regulation system. However, the interaction patterns between TF binding motifs remain elusive. The recent high-throughput assays, CAP-SELEX, have identified over 600 composite DNA sites (i.e. heterodimeric motifs) bound by cooperative TF pairs. However, there are over 25,000 inferentially effective heterodimeric TFs in human cell. It is not practically feasible to validate all heterodimeric motifs due to cost and labour. Therefore, it is highly demanding to develop a fast and accurate computational tool for heterodimeric motif synthesis.

**Results:** We introduce DeepMotifSyn, a deep-learning-based tool for synthesizing heterodimeric motifs from monomeric motif pairs. Specifically, DeepMotifSyn is composed of heterodimeric motif generator and evaluator. The generator is a U-Net-based neural network that can synthesize heterodimeric motifs from aligned motif pairs. The evaluator is a machine-learning-based model that can score the generated heterodimeric motif candidates based on the motif sequence features. Systematic evaluations on CAP-SELEX data illustrates that DeepMotifSyn significantly outperforms the current state-of-the-art predictors. In addition, DeepMotifSyn can synthesize multiple heterodimeric motifs with different orientation and spacing settings. Such a feature can address the shortcomings of previous models. We believe DeepMotifSyn is a more practical and reliable model than current predictors on heterodimeric motif synthesis.

**Availability and implementation:** The software is freely available at https://github.com/JasonLinjc/deepMotifSyn.

## 1 INTRODUCTION

Understanding the transcriptions factor (TF) recognition motifs is vital for the analysis of gene expression specificity [1, 2, 3, 4, 5, 6]. The related experimental technologies (e.g. ChIP-seq, ChIP-exo, and ChIP-nexus) have been developed to detect transcriptions factor binding sites (TFBSs) in various cell types and tissues, while some TFBSs do not have any strong-match motif [7, 8, 9, 10, 11]. Several studies have revealed that cooperative TF pairs can function as heterodimers for binding onto the composite DNA motif to regulate gene expression [12, 13, 14, 15, 16, 17]. Therefore, characterizing TF interactions and the corresponding heterodimeric DNA motifs are essential for understanding the function of non-coding DNA in gene regulation.

In a previous study, Jolma et al. developed CAP-SELEX (Consecutive Affinity-Purification Systematic Evolution of Ligands by Exponential Enrichment), to systematically investigated 9,400 TF pairs; and 315 of which were detected as heterodimeric DNA motifs [14]. That study deduced that there are around 25,000 effective heterodimeric TFs in human cell. However, it is not practically feasible to detect heterodimeric motifs of every potential TF pair owing to cost and labour. Wong et al. recently presented an machine-learning-based computational model, MotifKirin, to synthesize heterodimeric DNA motif [18]. They broke down the synthesis task into two phases: the phase one uses Random Forests to predict the orientation and overlapping length of two motifs; the phase two adapts a IOHMM (Input-Output Hidden Markov Model) to synthesize heterodimeric motifs based on the predicted orientation and overlap preference.

Despite such *in silico* modeling enables low-cost and rapid synthesis of heterodimeric DNA motifs in a DNA-binding family specific manner, it still has some shortcomings: 1) MotifKirin can only synthesize one heterodimeric motif for a specific monomeric motif pair, but practically, TF pairs often display more than one overlap and/or orientation case; 2) IOHMM was trained on the motifs for the same DNA-binding family, thus it is incapable of producing heterodimeric motifs from other family. Besides, one family only contains a few heterodimer motifs (12 per family averagely), such a few training data could hinder the generalizability of machine-learning-based models; 3) IOHMM performs badly on synthesizing motif pairs with long overlap than those with spacings; it reflects IOHMM has limitations on capturing complex patterns between two motifs. To address the above issues, we develop a deep-learning-based model to accurately synthesize heterodimeric DNA motifs across different DNA-binding families.

Deep learning has been making impressive advances in DNA motif research since Deep-Bind was proposed in 2015 [19]. DeepBind is the first to adapts convolutional neural network (CNN) to predict TFBSs from DNA or RNA sequences; it also introduced a subtle approach (i.e. mutation maps) to discover motifs from convolutional kernels [20]. Hassanzadeh et al. then proposed DeeperBind which uses a bi-directional long short-term memory (LSTM) network in addition to CNN. Their experiment showed DeeperBind surpassed DeepBind on the motif prediction task [21]. Other researchers have been unremittingly tapping the potential of various deep learning architectures to improve the accuracy, such as DanQ [22], DeepSEA [23], KEGRU [24], and iDeeps [25]. Most recently, Avsec et al. presented BPNet, a deep dilated convolutional neural networks with residual connections, predicts TF binding motifs at the base resolution [26]. Inspiringly, this study reported that a well-trained BPNet can identify the patterns of TF cooperativity (e.g. Oct4-Sox2) directly from DNA sequences. The studies above have demonstrated CNN is a feasible architecture for modeling TF binding patterns and identifying the corresponding DNA motifs.

However, unlike the previous tasks, heterodimeric motif synthesis aims at generating composite motif sequence patterns instead of simply predicting binary labels or assay profiles. In this work, we adapt a variant of CNN (i.e. U-Net [27]) to learn TF interactions and generate heterodimeric motifs. We then develop a machine-learning-based model to evaluate generative heterodimeric motifs. Together, we name our end-to-end motif synthesizer as DeepMotifSyn, which substantially outperforms the previous model as demonstrated through our experiments.

## 2 MATERIALS AND METHODS

### 2.1 Background of U-Net

U-Net is a symmetric convolutional network that was first designed for cellular image segmentation [27, 28, 29]. The general architecture of U-Net consists of three components: Contraction, Bottleneck, and Expansion. The contraction down-convolutes a image by extracting the spatial features of the image; the bottleneck compresses feature maps that preserves the most important information and reduces model complexity; the expansion module constructs a segmented image by up-convoluting compressed feature maps using transposed convolution operations. Remarkably, the most artful aspect of U-Net is skip connection, which concatenates the up-convoluting output in each expansion layer with the feature maps in the symmetric construction layer. The concatenated feature map is then propagated to the successive layer. Such structure enables the expansion module to retrieve spatial information lost in down-sampling, retaining the image integrity and mitigate the distortion when reconstructing the segmented image [30].

Comparing heterodimeric motif synthesis with image segmentation, we notice U-Net has great potentials on synthesizing composite motif sequences: 1) The CNN-based contraction of U-Net can be used as an encoder to extract TF-interaction patterns, particularly to learn how two monomeric motifs overlaps; 2) In addition to overlap, the flanking sequences should be well persevered during synthesizing. Skip-connection is ideally suited for network to retrieve non-overlapping sequences of two motifs when constructing the heterodimeric motif; 3) It has been shown that U-Net has impressive performance with small labeled training data [27]. Therefore, we designed a U-Net-based architecture to synthesize heterodimeric motifs from monemeric motif pairs.

### 2.2 DeepMotifSyn: proposed deep learning model

DeepMotifSyn consists of heterodimeric motif generator and evaluator. The generator is a U-Net-based neural network that down-convolutes a monomeric motif pair and then up-convolute to generate a heterodimeric motif. A downstream machine learning model is used as the evaluator to compute for the predicted probability that a generated heterodimeric motif is the true one, based on the motif sequence features and DNA-binding family. Together, the generator and evaluator provide an integrated tool that enables users to conveniently synthesize heterodimeric motifs using any motif pair of interests. Figure 1 illustrated how DeepMotifSyn can generate and score heterodimeric motifs from two motifs (i.e. ALX4 and EOMES): we first generated all possible alignments of two motifs based on four orientations [18] and up-to-19 spacing/overlap, each of which DeepMotifSyn generator synthesizes a specific heterodimeric motif. Then, DeepMotifSyn evaluator scores each generated heterodimeric motif for its possibility based on the motif sequence features (see Section 2.2.2).

**Figure 1:**
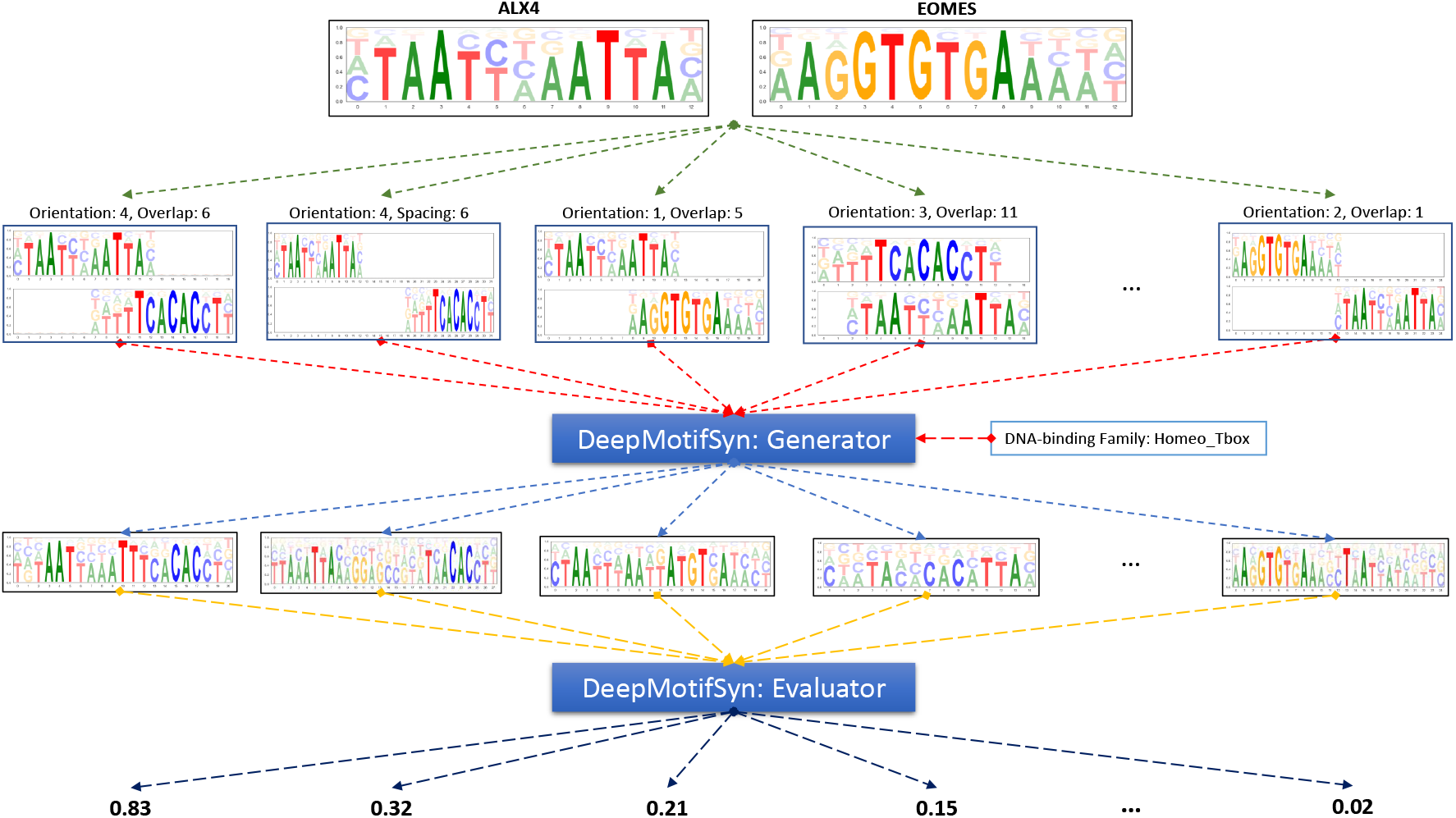
Illustrative example of how DeepMotifSyn synthesizes heterodimeric DNA motif. Given two monomeric motifs (e.g. ALX4 and EOMES), DeepMotifSyn first generates multiple heterodimeric motifs with all possible orientation and overlap/spacing cases. It then scores each candidate based on the motif sequence features. As observed from the CAP-SELEX data, top 10 generative heterodimeric motifs enough to be considered as Deep-MotifSyn’s final prediction candidates.

#### 2.2.1 Heterodimeric motif generator

The architecture of motif generator is illustrated in Figure 2. It contains a general U-Net backbone which involves contracting path, bottleneck module, an expansive path, and the subsequent convolutional layers with filter size of 1 produce the heterodimeric motif in the form of a position probability matrix (PPM). The input to the neural network was a 108-channel sequence involving a 8-channel (noting A, G, C and T of motif1 and motif2) motif pair and a 100-channel DNA-binding family one-hot encode.

**Figure 2:**
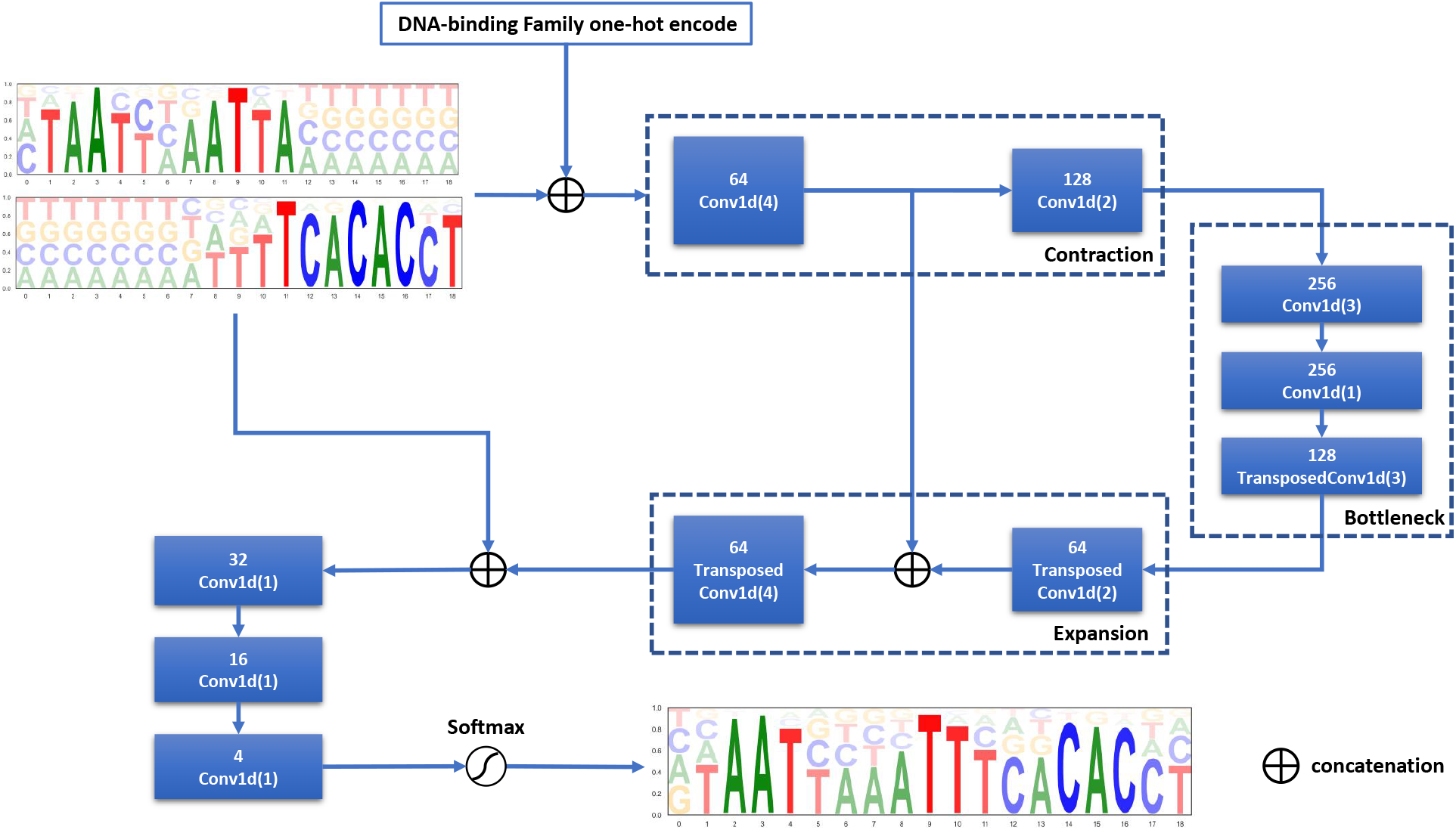
Architecture of the U-Net-based neural network that was trained to predict the Position Probability Matrix (PPM) of a heterodimeric motif from a aligned monemeric motif pair.

As Figure 2 shows, the contracting path down-samples input sequence by two convolutions with the filter length of 4 and 2 successively. The expansive path up-samples the previous feature maps along with the one in the symmetric contracting layer, using two transposed convolutions with the filter lengths of 2 and 4. In between, a bottleneck module compressed the feature maps using a 3-layer convolutional autoencoder. Lastly, the expansive feature maps are passed through a three-layers convolution network with filer with the length of 1, to predict the probability of nucleobases at each position. Note that each convolutional layer in this network was preceded by a batch-normalization and ReLU activation except the last layer. The last convolutional layer facilities a softmax operation to produce a position probability matrix.

As the heterodimeric motifs varied in sequence length, we designed a mask mean square loss as the cost function for network training. Specifically, we first pad every ground true heterodimeric motifs into a 35 × 4 matrix with zero vectors, mathematically noting as *M* = [*m*_1_,*m*_2_,…,*m*_35_],*m_i_* ∈ [0,1]^4^. 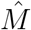 denotes the predictive heterodimeric motif accordingly. We defined a sign vector *δ* = [*a*_1_,*a*_2_,…, *α*_35_],*a_i_* ∈ {0,1}, to indicate the presence of non-zero elements in heterodimeric motif. The mask mean square error (MaskMSE) for the generated motif 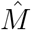 is defined as:

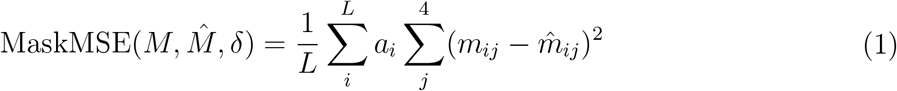

where 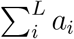 is the actual length of the heterodimeric motif, and *L* = 35 is our padded motif length. We then trained our network using Adam optimizer to minimize the MaskMSE loss, with the batch size of 100 for 200 epochs.

#### 2.2.2 Heterodimeric motif evaluator

Our evaluator is developed to predict the probability how much each heterodimeric motif candidate is the annotated one in gene regulation. To develop such a model, we carefully designed a 784-dimensional feature vector to represent the interaction of monomeric motif pairs. These features can be categorized into four types: motif pair sequence, generative motif sequence, motif pair orientation, motif pair overlapping, and DNA-binding family features.

Note that both the input motif pair and its generated motif are represented by the position probability matrix (PPM). To highlight the important position during modeling, we convert PPM into an information content matrix (ICM). Specifically, the total value of each position in PPM is scaled by information content (IC) which indicates the level of conservation [31, 32]. Mathematically, ICM at position *i* can be computed as:

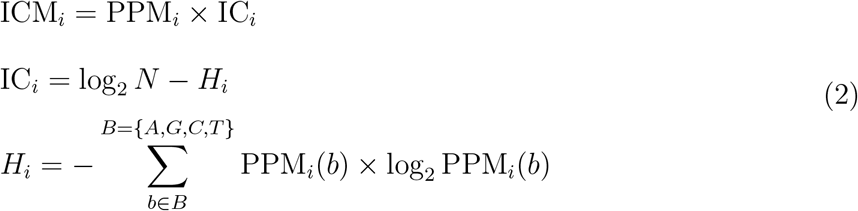

where *N* = 4 is the number of bases, *H_i_* denotes Shannon’s entropy [33] of PPM at position *i*, and PPM_*i*_(*b*) denotes the probability of base *b* appearing at position *i*. Figure 3 illustrates a comparative example of ICM and PPM. We then use the ICM and the positional entropy of each motif along with the generated heterodimeric motif as the sequence features.

**Figure 3:**
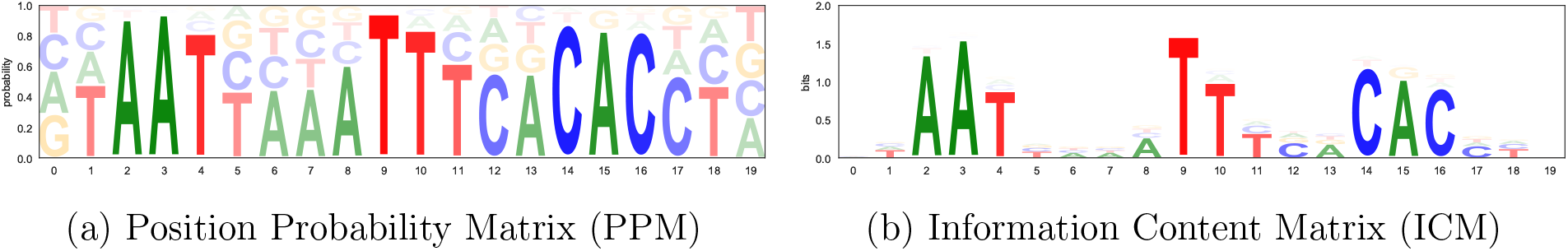
PPM and ICM of a heterodimeric DNA motif

Given a monomeric motif pair 〈*X, Y*〉 in double helix, there are four possible orientations to synthesize heterodimeric motif: 〈*X, Y*〉, 〈*Y, X*〉, 〈*y, X*〉 and 〈*X, y*〉, where *y* is the reversion complement of *Y* [18]. We encode the orientation case into a 4-bit one-hot vector. The motif pair alignment features can be categorized into spacing and overlapping. The prediction of overlap sequence is crucial to heterodimeric motif synthesis. We extract the ICMs and nucleobase entropy at every overlapping position as features. We also compute the Euclidean distance of two motifs’ ICM and entropy at the overlapping positions. We then sum the overlap ICM at position and nucleobase level respectively to store the iteration information. For those motif pairs without overlap, we use a zero vector as the placeholder during modeling. In addition, we use a numerical feature to represent the overlap/spacing length, of which the sign indicates the spacing (negtive) and overlap (positive). Lastly, we statistically analyze the orientation and overlapping length for each DNA-binding family, and build the features to represent the family-specific distribution of overlapping and orientation.

Based on the above carefully designed features, we have implemented and systemically evaluated five classifiers on selecting the actual heterodimeric motif. Our experiments below reveal that XGBoost with optimized hyper-parameters is promising in evaluating the generated heterodimeric motifs.

### 2.3 Data construction

The heterodimeric and monomeric motif dataset contains 614 heterodimeric motifs from 313 monomeric motif pairs detected by CAP-SELEX [14]. Each motif is represented by the position probability matrix. We padded every motif matrix with zero vectors into a 35 × 4 matrix. Note that the longest length of the heterodimeric motif in the dataset is 31. Thus, each motif pair can be represented by a 35 × 8 matrix as the input to neural network. In addition to 614 annotated motif pairs, we generated 368,381 possible alignments of 313 monomeric motif pairs to evaluate DeepMotifSyn, based on four orientation cases with the spacing/overlap length up to 19.

### 2.4 Evaluation Metrics

To evaluate DeepMotifSyn’s motif generator, we designed a motif position probability matrix (PPM) distance as the metric. Given a generated motif PPM 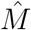 and a ground-true motif PPM *M*, we first align them using Needleman-Wunsch global profile alignment method [34]. Then, we can calculate the Euclidean distance at every aligned position:

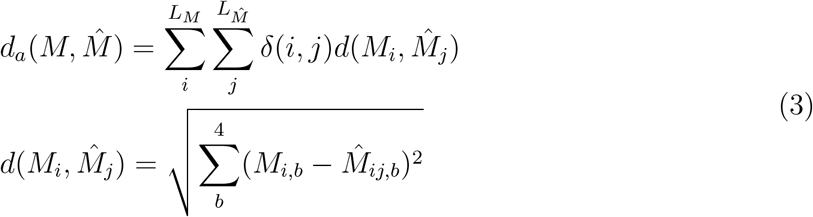

where *L* is the length of motif, and *δ*(*i,j*) = 1 indicates *M_i_* is aligned with 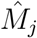, otherwise *δ*(*i,j*) = 0. We add the maximum distance of aligned base pairs 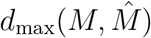 as the penalty to unaligned positions in the calculation of motif PPM distance 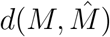:

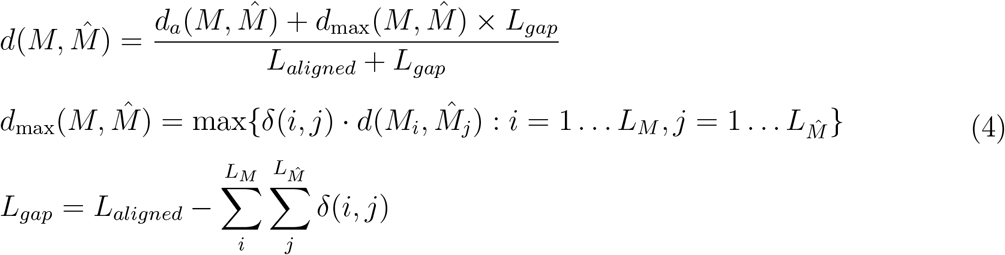

where *L_gap_* denotes the number of unaligned base pairs.

To estimate different machine learning models as the motif evaluator, we used precision-recall analysis instead of ROC (Receiver Operating Characteristic) analysis owing to the data imbalance issue. Each model candidate is optimized by a 10-fold cross-validated search over hyper-parameters settings. Lastly, we compared each optimized model on an hold-out testing dataset, and the model with the highest PR-AUC (Area under the Precision-Recall curve) will be used as our final motif evaluator.

## 3 RESULTS

Our experiments demonstrated DeepMotifSyn achieving promising performance on both generating and evaluating heterodimeric motif, outperforming the previous approach in the task of end-to-end heterodimeric motif synthesis.

### 3.1 Performance of DeepMotifSyn’s motif generator

To compare our study with previous studies fairly, we performed leave-one-motif-pair-out cross-validation to estimate our U-Net-based generator on 313 motif pairs which synthesize 614 heterodimeric motifs. Note that the previous IOHMM-based motif generator was trained on the motifs from one specific DNA-binding family. It is thus unable to handle the DNA-binding family with very few heterodimeric motifs. We first compared our U-Net-based network with IOHMM on 530 heterodimeric motifs from 45 DNA-Binding families. Table 2 shows U-Net outperformed IOHMM with an average motif PPM distance of 0.17. The difference between two predictions is statistically significant with a t-test p-value of 6.0 × 10-^44^. The bar chart in Figure 5 demonstrates that our U-Net-based generator significantly outperformed IOHMM across 40 DNA-binding families among 45, and the motif matrix distance error has been decreased averagely by 9% among 530 heterodimeric motif synthesis comparing to IOHMM-based motif generator. For the other 84 heterodimeric motifs from 42 DNA-Binding families, U-Net still achieved a comparative performance with an average motif PPM distance of 0.24, indicating that our network can leverage the motif interaction patterns across different families to synthesize heterodimeric motifs. Figure 4 demonstrates three heterodimeric motif synthesis examples, in which DeepMotifSyn showed its ability to predict how the individual motifs form composite DNA motif. The predicted sequence in the red box shows that our model has advantages over IOHMM on synthesizing overlapping motifs. Surprisingly, we found our model can predict the alteration of non-overlapping sites (see the sequence in the orange box), which reveals the complexity of genomics grammars and the feasibility of our U-Net-based network.

**Figure 4:**
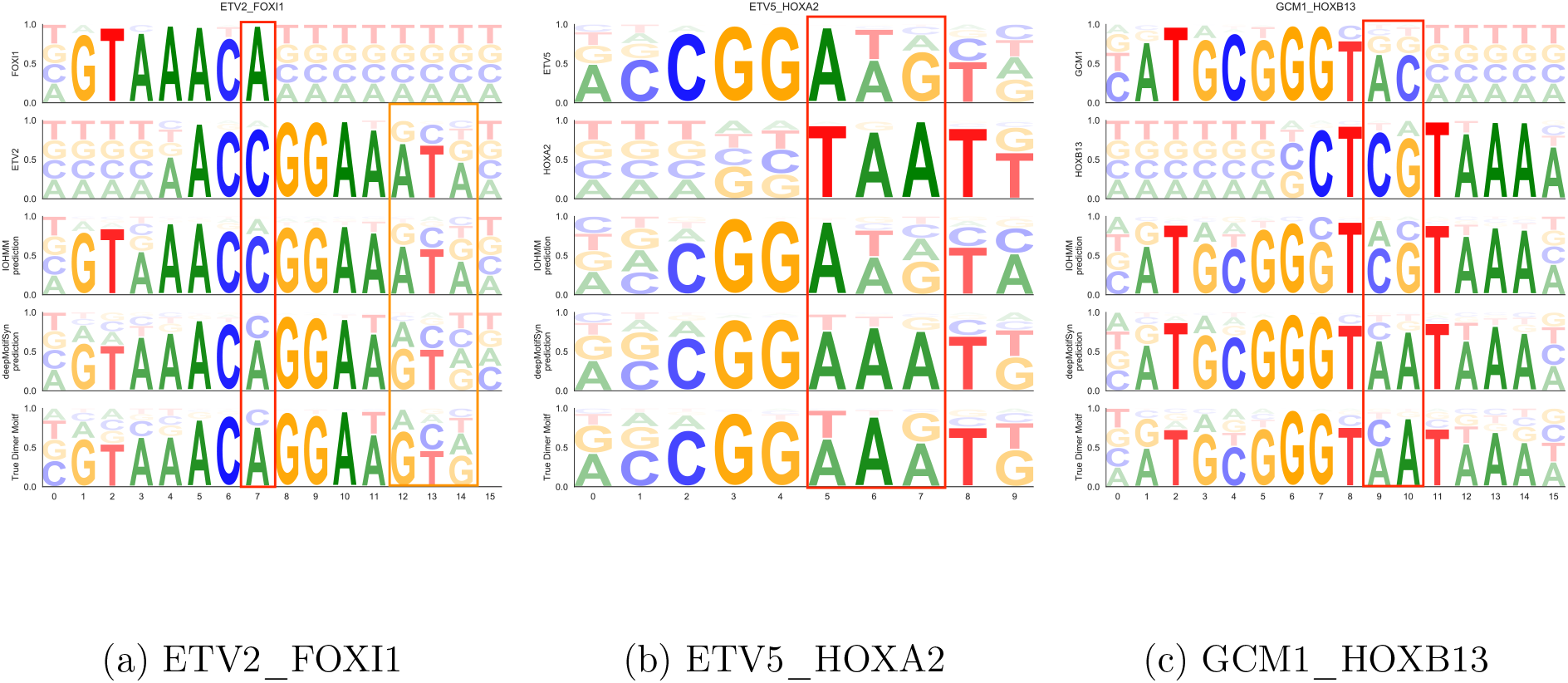
Comparison of IOHMM and U-Net-based neural network on heterodimeric motif synthesis with true orientation and overlap/spacing settings

**Figure 5:**
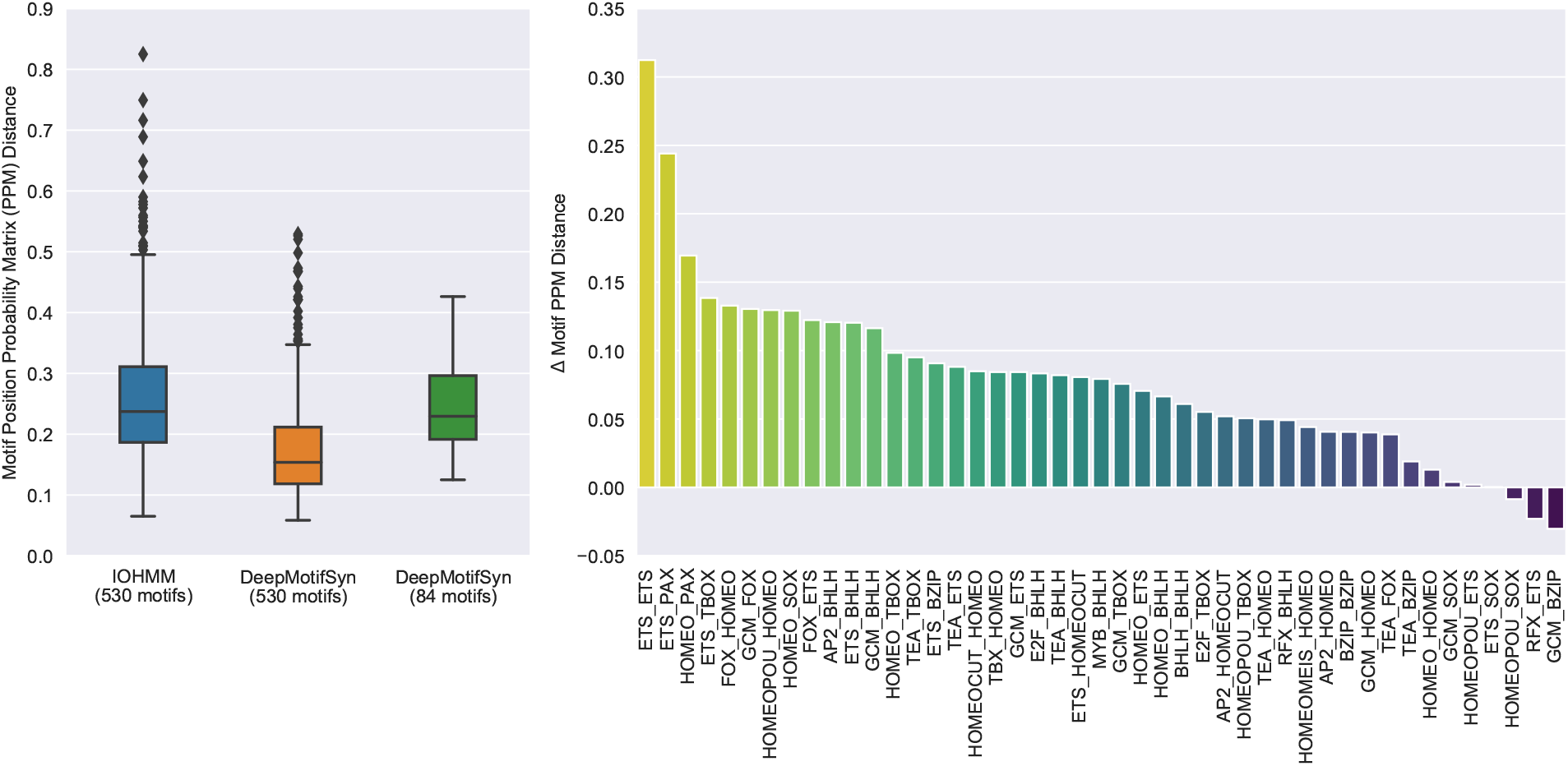
Performance of IOHMM and U-Net-based neural network under leave-one-motifpair-out cross-validation. The box-plot chart (left) shows the comparison of IOHMM and U-Net-based model on 530 heterodimeric motifs, and the performance of U-Net-based model on 84 motifs from small-sized DNA-binding families. The bar chart (right) demonstrates the average improvement of U-Net-based model on IOHMM across 45 DNA-binding families, where Δ motif PPM distance = PPM distance of IOHMM motif – PPM distance of U-Net-based motif.

**Table 1:** Comparison of U-Net-based neural network and IOHMM under leave-one-motifpair-out cross-validation

| Motif Generator | 530 heterodimers |  |  | 84 heterodimers |  |
| --- | --- | --- | --- | --- | --- |
|  | mean | std | p-value | mean | std |
| U-Net-based | 0.17 | 0.08 | $6.0 \times 10^{-44}$ | 0.24 | 0.07 |
| IOHMM-based | 0.26 | 0.11 |  | — | — |

**Table 2:** Comparison of DeepMotifSyn and MotifKirin under leave-one-motif-out crossvalidation on end-to-end heterodimeric motif synthesis

|  | 530 heterodimers |  |  | 84 heterodimers |  |
| --- | --- | --- | --- | --- | --- |
|  | mean | std | p-value | mean | std |
| DeepMotifSyn | 0.26 | 0.01 | $5.1 \times 10^{-43}$ | 0.28 | 0.08 |
| MotifKirin | 0.39 | 0.18 |  | — | — |

Our experiment demonstrated U-Net-based motif generator had better synthesis accuracy and generalization comparing to the previous IOHMM. Note that each IOHMM was developed for one specific DNA-binding family, it thus can only be evaluated on the motifs from the same family, of which the estimated performance can be easily overrated due to the limitation of validation data and the underlying over-fitting issue. By contrast, our model is more generalised as it was built on the motifs from multiple families. Besides, the estimated performance of our model under leave-one-out cross-validation on 313 motif pairs is much better than IOHMM’s inner-family cross-validation.

### 3.2 Performance of DeepMotifSyn’s motif evaluator

To find a suitable machine learning model as the motif evaluator, we tested five well-established model including histogram-based gradient boosting tree [35, 36], XGBoost [37], CatBoost [38], Random Forest [39], and extremely randomized trees [40] on 368,995 motif pairs. The whole dataset was divided into a training dataset (90%) and a testing dataset (10%). We then performed 10-fold cross-validated randomized search over hyper-parameter settings of five models on the training dataset, models with best hyper parameters were further compared on the independent testing dataset in terms of PR-AUC and ROC-AUC. Table 3 shows that XGBoost with optimized hyper-parameters achieved then best performance under both 10-fold cross-validation and independent testing among five models. XGBoost had significant advantages over the other four models based on precision-recall analysis with an average PR-AUC higher than 40%.

**Table 3:** Comparison of five machine learning models on the evaluation of heterodimeric motif candidates

| Model | 10-fold cross-validation | Testing |  |
| --- | --- | --- | --- |
|  | PR-AUC | ROC-AUC | PR-AUC |
| **XGBoost** | **0.405 ± 0.061** | **0.995** | **0.431** |
| Histogram Gradient Boosting Trees | 0.339 ± 0.059 | 0.994 | 0.396 |
| CatBoost | 0.365 ± 0.069 | 0.993 | 0.371 |
| Random Forest | 0.330 ± 0.051 | 0.975 | 0.337 |
| Extremely Randomized Trees | 0.284 ± 0.037 | 0.981 | 0.262 |

Moreover, we performed a leave-one-out cross-validation to further estimate XGBoost along with U-Net-based generator on 313 motif pairs. In each fold, we first generated all possible monomeric motif pairs based on 4 orientations and up-tp-19 overlapping/spacing, on which we applied our U-Net-based network to synthesize heterodimeric motif candidates. We then trained XGBoost on the handcrafted features of 312 motif pairs, and score the heterodimeric motif candidates derived from each leave-one-out motif pair. Note that the training motifs of DeepMotifSyn generator excludes the testing motif pair. In this 313-fold cross-validation, XGBoost achieved ROC-AUC of 0.99 and PR-AUC of 0.40 (supplementary), which was similar to its 10-fold cross-validation performance. Since the maximum heterodimeric motifs for a single motif pairs in our dataset is 10, we selected the 10 generated motifs with the highest scores as our final predictions. We observed such a selection strategy recovered 433 motifs (70%) among 614 CAP-SELEX-validated heterodimeric motifs. It’s also worth mentioning that top 30 XGBoost predictions of each motif pair covered 90% of ground-true motifs under leave-one-out cross-validation on 313 motif pairs.

Lastly, we compared our U-Net-based neural network together with XGBoost as DeepMotifSyn to MotifKirin on the end-to-end heterodimeric motifs synthesis task. Table 3 illustrated that DeepMotifSyn remarkably surpassed MotifKirin with a mean motif PFM distance of 0.36 on synthesizing 530 heterodimeric motifs. These 530 heterodimeric motifs are grouped by 45 DNA-binding families, as Figure 6 shows, DeepMotifSyn has significantly better performance on the 41 families. The lower standard deviation also indicates DeepMotifSyn is more robust than MotifKirin. Due to the limitation of IOHMM model, MotifKirin can not synthesize the other 84 heterodimeric motifs from small-sized families (which contains no more than two heterodimeric motifs). On the contrary, deepMotifSyn demonstrated its impressive capability in handling such motifs, achieving an average motif PPM distance of 0.28 on 84 heterodimeric motifs.

**Figure 6:**
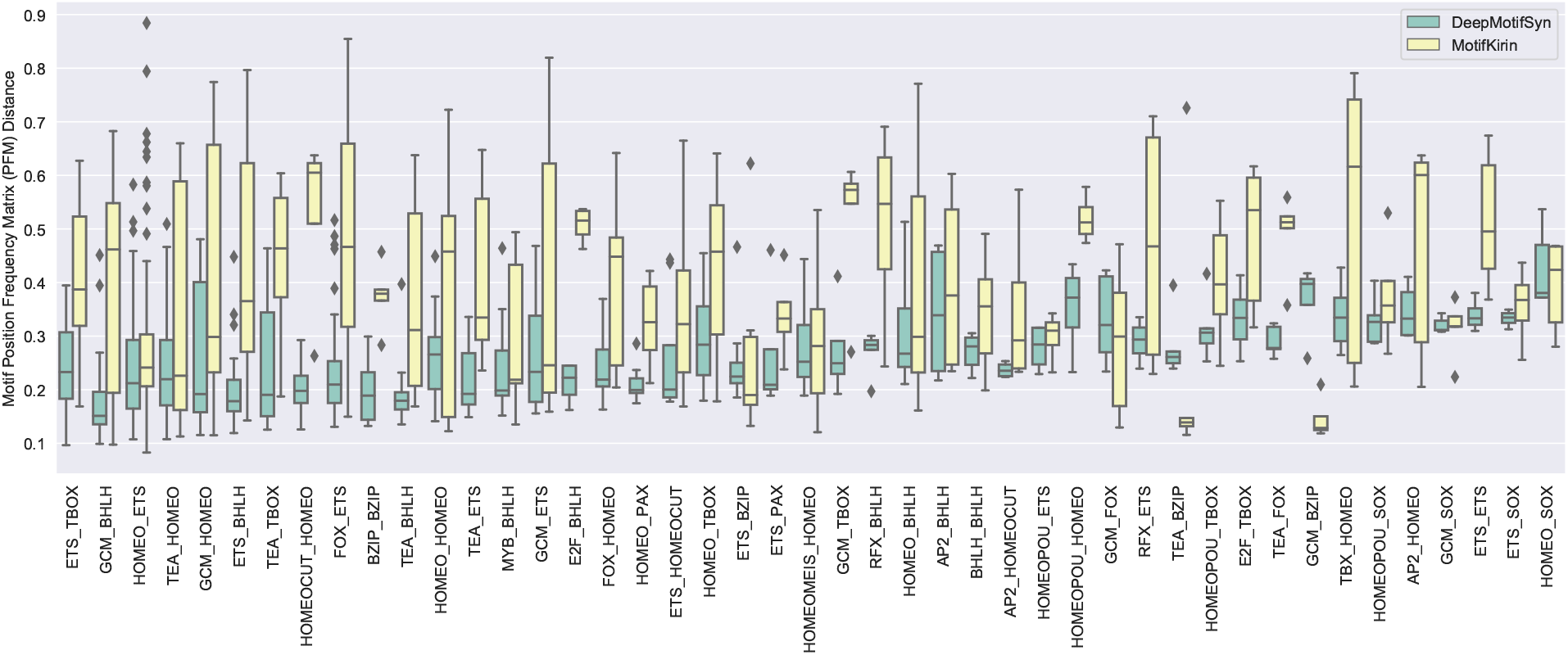
Comparison of DeepMotifSyn and MotifKirin under leave-one-motif-out crossvalidation across 45 DNA-binding families. The experimental settings are the same as our previous study [18].

### 3.3 Application of DeepMotifSyn on end-to-end heterodimeric motif synthesis

Herein, we demonstrated an example of how DeepMotifSyn synthesizes FLI1-FOXI1 heterodimeric motif. Note that this motif is independent of our training data. As Figure 7 shows, there are four heterodimeric motifs validated by CAP-SELEX derived from two monomeric motifs, FLI1 and FOXI1. Taking a position probability matrix pair as the input, DeepMotifSyn generated 1,218 heterodimeric motif candidates based on four possible orientations with the up-to-7 overlapping and up-to-13 spacing. The longest overlapping is the length of the short monomeric motif. We set 35 as the maximum length of generative heterodimeric motif for our model. Each generated motif is attached with a score predicted by DeepMotiSyn evaluator. Figure 7 illustrated the top 3 candidates were well matched with the validated heterodimeric motifs. The candidates with the highest deepSynMotif score has the lowest motif PPM distance with FLI1-FOXI1-1 among 1,218 generative motifs. The best matched generative motif of FLI1-FOXI1-4 ranks 41 among the candidates. We also applied MotiKirin to synthesize the FLI1-FOX1 motif, it can only produce one motif which is better aligned with FLI1-FOXI1-3 than the others. Nevertheless, DeepMotifSyn’s synthesis of FLI1-FOXI1-3 significantly surpassed MotifKirin with 27% improvement on motif PFM distance. Interestingly, we noticed the DeepMotifSyn score somehow represents the quality of the candidate even if we trained our model in a classification manner. In general, this case study demonstrated our DeepMotifSyn is a more practical and accurate approach for heterodimeric motif synthesis comparing to MotifKirin.

**Figure 7:**
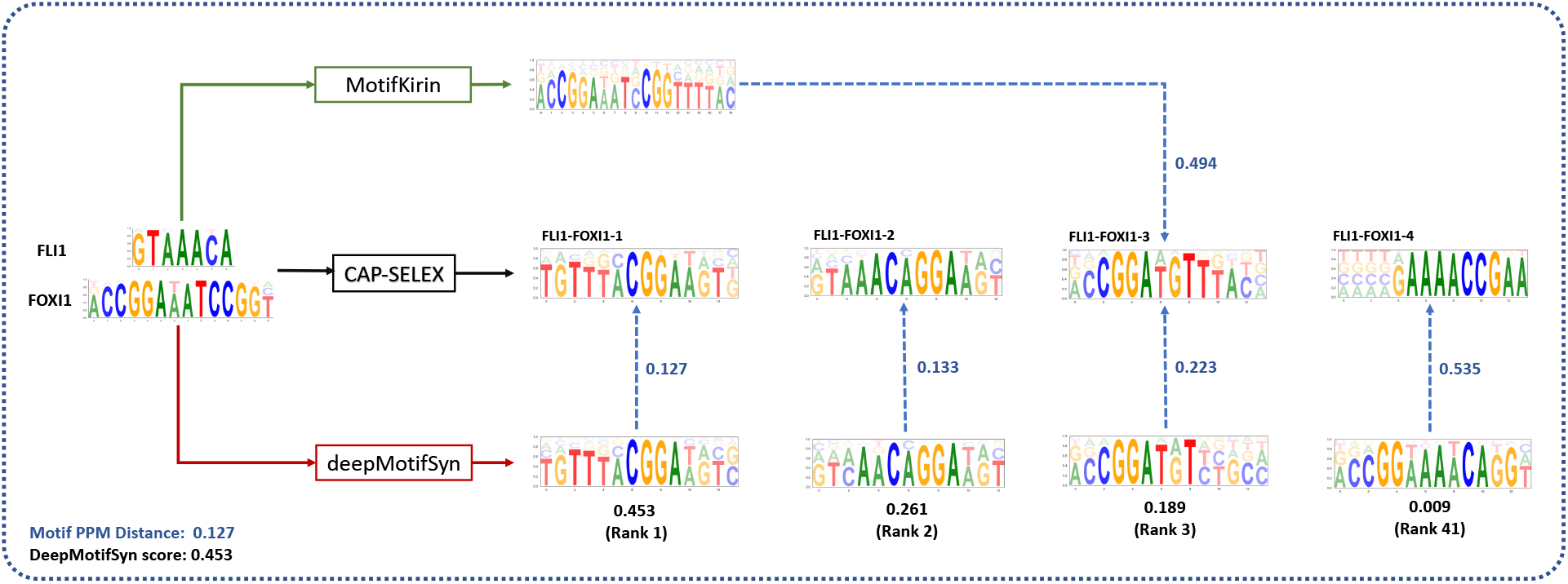
Heterodimeric motif synthesis on FLI1 and FOXI1 using DeepMotifSyn and MotifKirin. Note that MotifKirin can only synthesize one heterodimeric motif from a monomeric motif pair. The value in blue is the motif PPM distance of predictive and true heterodimeric motif, the value in black is DeepMotifSyn score.

## 4 Discussion

In this work, we introduced a deep-learning-based approach to synthesize heterodimeric motifs from monomeric motif pairs. Through systematic investigation, we illustrated that our newly developed suite of models, DeepMotifSyn, outperforms the current state-of-the-art method with a transformative way on synthesizing heterodimeric motifs. The previous model generates the heterodimeric motif based on separated predictive spacing length and orientation of the monomeric motif pair. Such a synthesis approach has some substantial limitations, since most of TF-TF pairs’ cooperatively bound sits involves more than one orientation and various spacing preferences [14, 16]. By contrast, DeepMotifSyn generates and scores heterodimeric motif candidates with all potential orientations and spacing preferences (see Figure 1). Our experiment demonstrated that DeepMotifSyn-synthesized heterodimeric motif candidates were able to recover 70% of *bona fide* heterodimeric motifs validated by CAP-SELEX. In addition, we systematically evaluated MotifKirin on synthesizing heterodimeric motifs given ground true orientation and spacing settings, showing that DeepMotifSyn significantly surpassed MotifKirin on the motif synthesis of 40 heterodimer DNA-binding families. DeepMotifSyn also leverages the motifs of multiple DNA-binding families to synthesize the heterodimeric motif for new family, which is a substantial feature MotifKirin lacks.

We expect our DeepMotifSyn can be improved through training on additional heterodimeric motif datasets subject to its availability. We also envision our deep-learningbased model can be applied to hetero-multimeric motif synthesis by taking multiple motifs as input in the future.

## Acknowledgment

The work described in this paper was substantially supported by the grant from the Research Grants Council of the Hong Kong Special Administrative Region [CityU 11200218], one grant from the Health and Medical Research Fund, the Food and Health Bureau, The Government of the Hong Kong Special Administrative Region [07181426], and the funding from Hong Kong Institute for Data Science (HKIDS) at City University of Hong Kong. The work described in this paper was partially supported by two grants from City University of Hong Kong (CityU 11202219, CityU 11203520). This research was substantially sponsored by the research project (Grant No. 32000464) supported by the National Natural Science Foundation of China and was substantially supported by the Shenzhen Research Institute, City University of Hong Kong. We gratefully acknowledge the support of NVIDIA Corporation with the Titan XP GPU for this research.

